# Copeptin response to acute physical stress during hypohydration: An exploratory secondary analysis

**DOI:** 10.1101/2021.09.19.460958

**Authors:** Harriet A. Carroll, Olle Melander

## Abstract

**Purpose:** Plasma copeptin (a surrogate marker of arginine vasopressin) is increasingly being used as a marker of stress in several research and clinical contexts. However, the response to an acute physical stressor in healthy adults has not yet been tested meaning it is unclear whether copeptin reflects dynamic changes in stress or whether this is moderated by different basal copeptin concentrations and how this relates to other stress hormones.

**Methods:** Secondary data analysis in a subsample of participants (n = 7; n = 1 woman) who opted-in for muscle biopsies in a randomised crossover study investigating the effects of acute hypohydration (HYPO) *versus* rehydration (RE) on metabolism.

**Results:** Plasma copeptin responded to the muscle biopsy stress stimulus, with a similar magnitude of difference according to basal concentrations during HYPO and RE; however, the peak was higher and concentrations typically took longer to return to baseline during HYPO. Despite large differences in copeptin concentrations, adrenocorticotropin hormone (ACTH) and cortisol showed a similar trends in response to the biopsies, regardless of hydration status.

**Conclusion:** Copeptin responded dynamically to an acute physical stressor (muscle biopsy). HYPO induced higher basal copeptin concentrations compared to RE, and resulted in a higher and prolonged copeptin response. Further research should investigate the mechanisms underlying the lack of differences in ACTH and cortisol according to hydration status.

## Introduction

Arginine vasopressin (AVP) is a hormone typically associated with fluid balance. AVP is also implicated in the hypothalamic-pituitary-adrenal (HPA) axis, though its exact role in this capacity in humans is unclear (Aguilera, 1998, Goncharova, 2013). AVP works synergistically with corticotropin-releasing hormone (CRH) (Goncharova, 2013). CRH is a strong secretagogue of adrenocorticotropic hormone (ACTH), and ultimately leads to cortisol secretion (Gibbs, 1986, Herman *et al*., 2016).

Whilst several animal models have attempted to characterise the AVP stress response (e.g. Lolait *et al*., 2007, Ma, Levy & Lightman, 1997), there is a paucity of causal human research (Goncharova, 2013). The limited evidence from humans focuses on stressors such as exercise (e.g. Negrao *et al*., 2000), acute cardiac events (e.g. Smaradottir *et al*., 2017), or surgery (e.g. Jochberger *et al*., 2009) which are multisystem, stress-inducing, and extreme events. Psychological stress has also been examined, demonstrating < 2 pmol·L^−1^ increase in copeptin concentrations (Siegenthaler *et al*., 2014, Urwyler *et al*., 2015); comparatively, our recent work has demonstrated hypohydration can increase copeptin by an average of 14.4 pmol·L^−1^ without having an effect on ACTH or cortisol (Carroll *et al*. 2019). Whilst it is well established that acute stress raises other stress hormones (such as cortisol), the role or time trend of the AVP response in this pathway is currently unknown, and its role in metabolic health is debated (Carroll & James, 2019).

Further, having a short half-life, among other factors, makes AVP difficult to measure (Hofland, Bakker & Feelders, 2015, Szinnai *et al*., 2007). Copeptin (the C-terminal part of the prepro-AVP hormone) has therefore been used as surrogate marker for AVP secretion, with its use being analogous to that of C-peptide for insulin (Morgenthaler *et al*., 2008). The responsiveness (i.e. how dynamically it changes in plasma measures in response to a stimulus such as acute stress) of circulating copeptin is also unclear due to its relative stability and longer half-life compared to AVP (Beglinger, Drew & Christ-Crain, 2017).

We recently conducted a randomised crossover trial investigating the acute effect of hydration status on the glycaemic response (Carroll *et al*., 2019). As part of the study, seven (out of 16) participants agreed to have muscle biopsies taken before having an oral glucose tolerance test (OGTT) in both a hypohydrated (HYPO) and rehydrated (RE) state. Thus we have obtained a unique dataset in which the acute response to an inadvertently stressful physical stimulus was measured, in a state whereby AVP (as measured by copeptin) is basally high (HYPO), or low (RE). The aim of this paper is to explore how acute physical stress affects copeptin and other stress hormones (ACTH and cortisol), and explore how this interacts with glycaemia as a marker of metabolic health.

## Methods

### Participants

Sixteen healthy adult volunteers (n = 8 men) participated in the main study (Carroll *et al*., 2019). Of these 16, nine participants agreed to have muscle biopsies, with a total of 7 participants having biopsies in both the rehydrated and hypohydrated trial arms. We did not obtain pre-biopsy blood samples from one participant, leaving a total sample in these analyses of n = 6. Baseline participant characteristics of the six volunteers who had both muscle biopsies suggests participants were in a similar physiological state before the intervention, except lower serum osmolality before the RE intervention (**Table 1**).

**Table 1.** Baseline participant characteristics (median, interquartile range) in a fasted and euhydrated state before the hydration interventions (n = 6)

|  | HYPO | RE | P-value |
| --- | --- | --- | --- |
| Sex (n male) | 5 | - | - |
| Age (y) | 31 (24, 40) | - | - |
| Randomisation (n first) | 4 | 3 | - |
| Body mass index (kg/m<sup>2</sup>) | 24.2 (22.5, 27.4) | 24.0 (22.1, 27.0) | 0.463 |
| Body mass (kg) | 73.5 (70.3, 76.9) | 75.6 (69.0, 79.1) | 0.463 |
| Urine specific gravity | 1.015 (1.013, 1.017) | 1.013 (1.012, 1.014) | 0.102 |
| Urine osmolality (mOsm·kg<sup>-1</sup>) | 445 (391, 492) | 415 (377, 450) | 0.249 |
| CSMA (mm<sup>2</sup>) | 13582(11413, 15908) | 13673 (11467, 15635) | 0.249 |
| Serum osmolality (mOsm·kg<sup>-1</sup>) | 287 (284, 293) | 284 (283, 286) | 0.042 |
| Plasma copeptin (pmol·L<sup>-1</sup>) | 4.63 (3.10, 6.00) | 4.77 (3.45, 5.40) | 0.753 |
| Plasma ACTH (pmol·L<sup>-1</sup>) | 3.23 (2.83, 3.43) | 3.88 (2.12, 4.76) | 0.917 |
| Plasma cortisol (nmol·L<sup>-1</sup>) | 281 (201, 373) | 229 (185, 351) | 0.345 |
| Serum glucose (mmol·L<sup>-1</sup>) | 5.18 (4.98, 5.37) | 5.02 (4.81, 5.35) | 0.345 |
| Serum insulin (pmol·L<sup>-1</sup>) | 27.03 (21.96, 40.02) | 23.59 (19.16, 34.52) | 0.893 |

Statistical significance calculated using Wilcoxon signed rank test. Abbreviations: ACTH, adrenocorticotropin hormone; CSMA, cross-sectional muscle area; HYPO, hypohydration trial arm; RE, rehydration trial arm

### Protocol

Full details of the study protocol have been published previously (Carroll *et al*., 2019). Briefly, in this randomised crossover trial, participants matched their diet (weighed and replicated food diaries) and physical activity energy expenditure (ActiHeart™) for three days prior to starting each intervention. On the third day of the pre-trial control phase, participants consumed ≥ 40 mL/kg lean body mass of non-alcoholic fluid to ensure euhydration prior to starting the intervention. The following day, participants came to the laboratory in a food and fluid restricted state from 2200 h the previous night (herein referred to as ‘euhydrated baseline’), had a peripheral quantitative computer tomography (pQCT) scan of the midpoint of their right thigh, a fasted blood sample, and then were put in a sweat suit in a 45 ± 2 °C heat tent for one hour, with pre- and post-heat tent body mass measurements. After the heat tent, participants chose a sandwich to consume containing ≥ 1 g salt. Subsequently, they were only allowed to consume low-water content foods, matched in both trial arms.

After the dehydration protocol in the heat tent, in the RE trial arm, participants were provided with 40 mL/kg lean body mass + 150 % sweat losses of water to consume until 2200 h, metered throughout the day. In the HYPO trial arm, participants were provided with 3 mL/kg body mass of water to consume until 2200 h. No other fluid was allowed to be consumed in both trial arms. With the exception of water, diet (low water content foods only) and physical activity were replicated on both trials after the heat tent.

The following day (herein the ‘full trial day’), participants came back to the laboratory and had a second fasted pQCT scan and blood sample. For those who had opted in, a muscle biopsy was taken in the fasted state from the vasterus lateralis muscle (Bergström, 1975), in a quasi-randomised manner for dominant or non-dominant leg first (pre-OGTT). A post-biopsy fasted blood sample was then drawn, before commencing a 2-hour oral glucose tolerance test (75 g glucose; Polycal, Nutricia, England), with bloods drawn every 15 min (**Figure 1**). After two hours, opted-in participants had a second muscle biopsy which is not relevant for these analyses.

**Figure 1.**
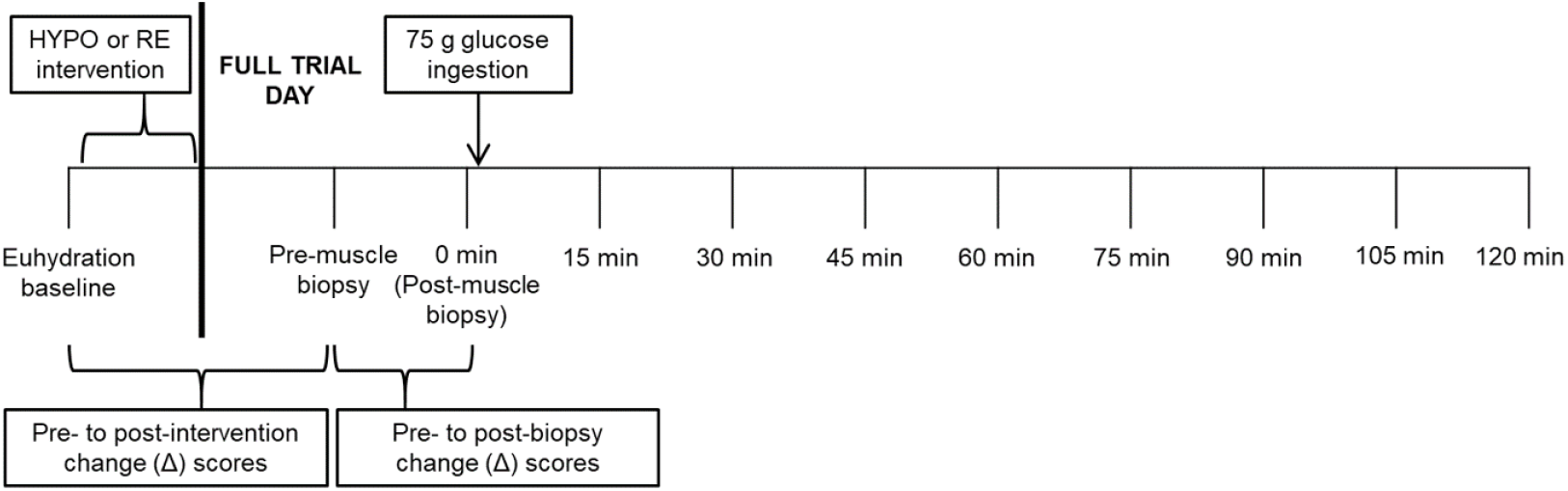
Schematic of timepoints of blood samples used in this sub-analysis. After the euhydration baseline sample, participants underwent a dehydration protocol (heat-tent) followed by rehydration (RE) or fluid restriction (HYPO), in a randomised crossover fashion

### Biochemical analysis

EDTA plasma was used to measure ACTH and cortisol via electrochemiluminescence immunoassays (Roche), and copeptin via an automated immune analyser (ThermoFisher Kryptor Compact Plus). Serum glucose was measured via a spectrophotometric assay (RX Daytona, Randox Laboratories), and serum insulin with a commercially available ELISA (Mercodia). Freezing-point depression was used to determine serum osmolality (Gonotex Osmomat auto) and urine osmolality (Micro-Osmometer 3300), and urine specific gravity was measured with a handheld refractometer (SUR-NE Clinical Refractometer; Atago). Further details can be found in Carroll *et al*. (2019).

### Data analysis

Descriptive statistics are shown as median (interquartile range) and analysed using Wilcoxon rank test. To determine the difference in analytes from euhydrated baseline to post-intervention (i.e. HYPO or RE), and the difference between pre- to post-biopsy, a change score (Δ) was calculated by subtracting the latter value from the former value, with accompanying 95 % confidence intervals (CI) of the change. In participant 01, we did not obtain a pre-biopsy blood sample in either trial; as such, these data have been excluded from all analyses (final sample n = 6). In order to understand the relationship between Δ scores, curve estimates were run, and either simple linear regressions were conducted to obtain *r*^*2*^ values, or non-linear (quadratic) pseudo-*r*^*2*^ values were obtained, depending on the data distribution.

It should be noted that the original study was powered to detect a change in the glycaemic response (n = 16) based on the expected response from pilot work (Carroll *et al*., 2019). Thus any inferential statistics in this manuscript are underpowered, and are presented in order to give the reader an understanding of where effects may be (i.e. they are provided in order to aid hypothesis development due to the exploratory nature of the analysis, rather than to provide definitive statistic inferences). Data were analysed on Excel (Microsoft Corp) and SPSS (IBM, version 26).

Time points in figures relate to the OGTT administration, thus time point “15 min” represents approximately 30 minutes post-biopsy. A schematic of the timepoints of the blood samples discussed in this sub-analysis can be found in **Figure 1**. All participants gave informed consent before participating (ethical approval reference number: 16/SW/0057; the original trial was registered at Clinicaltrials.gov: NCT02841449 and Open Science Framework: https://osf.io/ptq7m). Data are available from: doi: 10.15125/BATH-00547.

## Results

### Euhydrated baseline to post-intervention change (Δ) in participants who underwent muscle biopsies

In this sub-sample of six participants who had muscle biopsies on both trial arms, urine specific gravity and urine osmolality confirmed compliance to the protocol, along with Δ −1.9 (95 % confidence interval [CI] −3.1, −0.7) kg (∼2.6 %) body mass loss after HYPO compared to relative weight stability (Δ +0.07 [95 % CI −0.2, 0.3 kg [∼0.1 %] increased body mass) after RE (**Table 2**). Further, serum osmolality increased by 9 (95 % CI 8, 11) mOsm·kg^−1^ after HYPO with an accompanying average increase in plasma copeptin concentration of 17.49 (95 % CI 4.39, 30.60) pmol·L^−1^ (**Table 2**). These changes in serum osmolality and plasma copeptin concentration were not found after RE (average increases of 2 mOsm·kg^−1^, and 1.28 pmol·L^−1^, respectively). Plasma ACTH and cortisol, serum glucose and insulin concentrations remained stable from fasted euhydrated baseline to fasted full trial pre-biopsy, after both HYPO and RE (**Table 2**).

**Table 2.** Median (interquartile range) of markers of hydration status for HYPO and RE, and mean change (Δ, 95 % confidence interval) from euhydrated baseline to HYPO and RE pre-biopsy full trial day (n = 6)

| | Hypohydration | | Rehydration | | $p_{\text{difference full trial}}$<br>$\Delta$ HYPO vs. $\Delta$ RE | $p_{\text{difference full trial}}$<br>HYPO vs. RE |
| --- | --- | --- | --- | --- | --- | --- |
| | Full trial (pre-biopsy) | $\Delta$ baseline to full trial | Full trial (pre-biopsy) | $\Delta$ baseline to full trial | | |
| Body mass (kg) | 71.2 (66.9, 75.7) | -1.9 (-3.1, -0.7) | 72.4 (69.0, 77.6) | 0.07 (-0.2, 0.3) | 0.028 | 0.028 |
| CSMA (mm<sup>2</sup>) | 13425 (11127, 15844) | -446 (-1146, 254) | 13476 (11539, 15987) | 100 (-285, 485) | 0.116 | 0.116 |
| Urine specific gravity | 1.028 (1.026, 1.028) | 0.012 (0.006, 0.017) | 1.017 (1.017, 1.019) | 0.005 (0.003, 0.008) | 0.046 | 0.027 |
| Urine osmolality (mOsm·kg<sup>-1</sup>) | 939 (905, 980) | 454 (294, 615) | 566 (539, 622) | 199 (100, 297) | 0.046 | 0.028 |
| Serum osmolality (mOsm·kg<sup>-1</sup>) | 297 (294, 303) | 9 (8, 11) | 287 (285, 290) | 2 (-2, 6) | 0.027 | 0.027 |
| Plasma copeptin (pmol·L<sup>-1</sup>)<sup>a</sup> | 22.83 (12.58, 28.79) | 17.49 (4.39, 30.60) | 4.72 (3.88, 6.26) | 1.28 (-2.54, 5.11) | 0.043 | 0.043 |
| Plasma ACTH (pmol·L<sup>-1</sup>) | 2.21 (1.73, 2.81) | -1.29 (-2.83, 0.25) | 2.72 (2.25, 3.76) | -0.63 (-1.66, 0.41) | 0.173 | 0.075 |
| Plasma cortisol (nmol·L<sup>-1</sup>) | 299 (256, 325) | 14 (-69, 98) | 275 (227, 315) | 14 (-38, 66) | 0.917 | 0.463 |
| Serum glucose (mmol·L<sup>-1</sup>) | 5.27 (5.23, 5.41) | 0.09 (-0.29, 0.47) | 5.11 (5.01, 5.29) | -0.01 (-0.28, 0.26) | 0.345 | 0.249 |
| Serum insulin (pmol·L<sup>-1</sup>) | 24.14 (21.88, 26.97) | -5.54 (-17.29, 6.21) | 20.46 (18.59, 21.20) | -6.39 (-16.61, 3.84) | 0.500 | 0.043 |

<sup>a</sup> n = 5 for copeptin HYPO due to a missing sample. Statistical significance was calculated using paired Wilcoxon signed rank test. Abbreviations: ACTH, adrenocorticotropin hormone; CSMA, cross-sectional muscle area; HYPO, hypohydration trial arm; RE, rehydration trial arm

### Pre- to post-biopsy changes

There were no changes in serum osmolality or serum insulin concentration in either trial arm (**Table 3**). Both plasma ACTH (HYPO Δ 6.30, 95 % CI 1.91, 10.70 pmol·L^−1^; RE Δ 6.23, 95 % CI 0.69, 11.77 pmol·L^−1^) and cortisol (HYPO Δ 119, 95 % CI 23, 214 nmol·L^−1^; RE Δ 109, 95 % CI 52, 165 nmol·L^−1^) concentrations increased after the biopsy, by comparable amounts in both HYPO and RE trials (**Table 3**).

**Table 3.** Mean change (95 % confidence interval) pre- and post-biopsy of plasma/serum markers (n = 6)

| | Hypohydration | | Rehydration | | $p_{\text{difference full trial } \Delta \text{HYPO vs. } \Delta \text{RE}}$ | $p_{\text{difference post-biopsy HYPO vs. RE}}$ |
| --- | --- | --- | --- | --- | --- | --- |
| | Post-biopsy | $\Delta$ pre- to post-biopsy | Post-biopsy | $\Delta$ pre- to post-biopsy | | |
| Serum osmolality (mOsm·kg<sup>-1</sup>) | 298 (291, 303) | 1 (-3, 1) | 287 (239, 289) | 0 (-2, 2) | 0.336 | 0.002 |
| Plasma copeptin (pmol·L<sup>-1</sup>) | 55.98 (-2.22, 114.18) | 37.14 (2.34, 71.95) | 17.44 (-3.86, 38.74) | 11.59 (-1.97, 25.15) | 0.080 | 0.052 |
| Plasma ACTH (pmol·L<sup>-1</sup>) | 8.62 (2.66, 14.58) | 6.30 (1.91, 10.70) | 9.18 (0.83, 17.53) | 6.23 (0.69, 11.77) | 0.753 | 0.826 |
| Plasma cortisol (nmol·L<sup>-1</sup>) | 419 (289, 548) | 119 (23, 214) | 386 (275, 496) | 109 (52, 165) | 0.893 | 0.418 |
| Serum glucose (mmol·L<sup>-1</sup>) | 5.69 (5.18, 6.21) | 0.37 (0.07, 0.68) | 5.32 (4.82, 5.180) | 0.15 (-0.004, 0.31) | 0.138 | 0.080 |
| Serum insulin (pmol·L<sup>-1</sup>) | 31.41 (18.77, 44.05) | 6.67 (-1.91, 15.25) | 30.46 (19.65, 41.27) | 9.19 (-0.56, 18.94) | 0.893 | 0.872 |

Statistical significance calculated using paired Wilcoxon signed rank test. Abbreviations: ACTH, adrenocorticotropin hormone; HYPO, hypohydration trial arm; RE, rehydration trial arm

After both HYPO and RE, serum insulin tended to increase post-biopsy by comparable amounts (**Table 3**). In contrast, serum glucose did not change pre- to post-biopsy during RE, but did increase slightly during HYPO (Δ 0.37, 95 % CI 0.07, 0.68 mmol·L^−1^). Post-biopsy, plasma copeptin increased after HYPO (Δ 37.17, 95 % CI 2.34, 71.95 pmol·L^−1^), and less reliably in RE (Δ 11.59, 95 % CI −1.97, 25.15 pmol·L^−1^) (**Figure 2**). These changes in serum glucose and plasma copeptin during HYPO were not significantly different to the changes during RE (glucose *p* = 0.138; copeptin *p* = 0.080; **Table 3**). Further, HYPO exaggerated the copeptin response to the biopsies compared to RE (Supplementary Information, and Figure S1).

**Figure 2.**
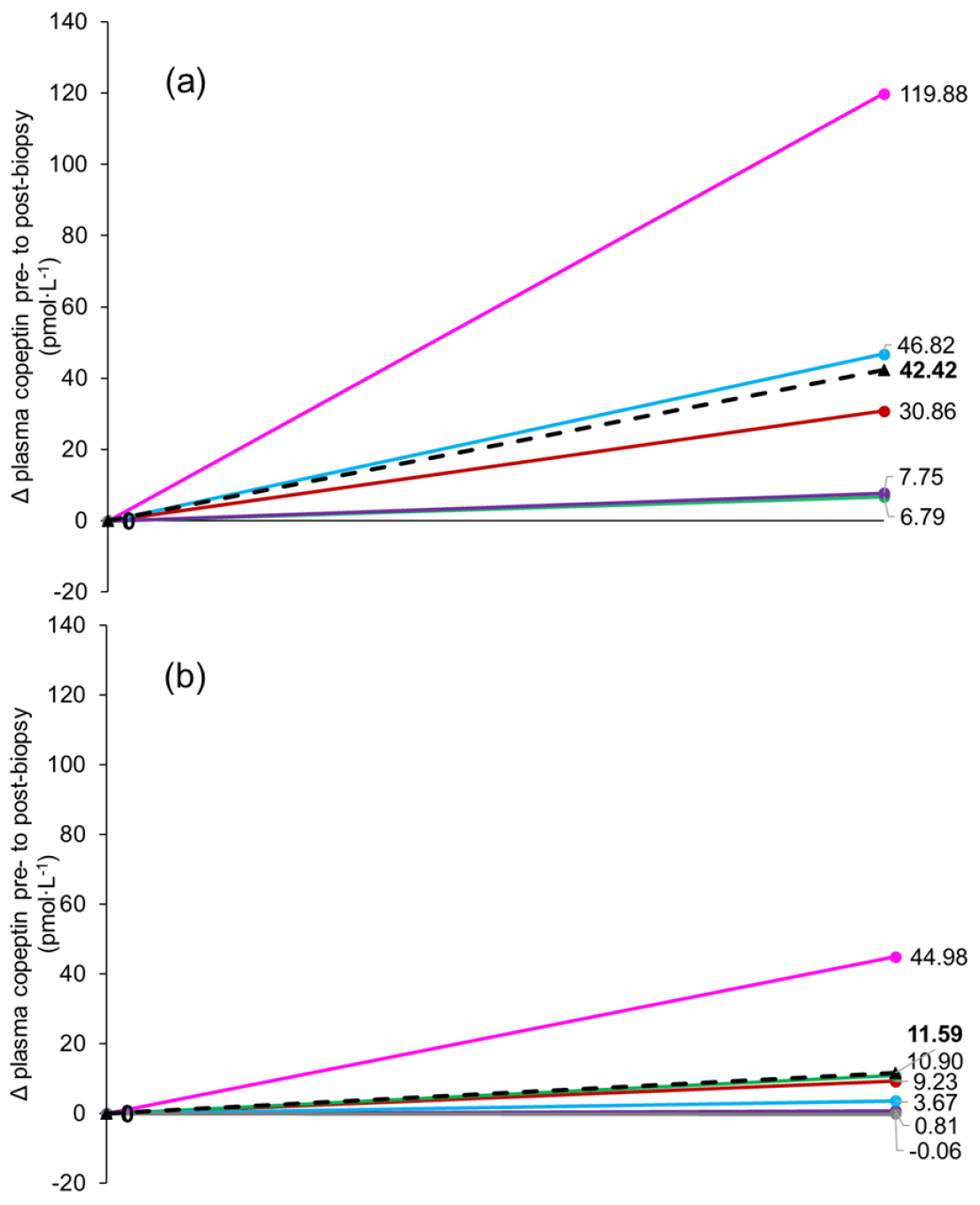
Individual (coloured lines) and average (black dashed lines) Δ plasma copeptin concentration pre- to post-biopsy in the (a) hypohydrated (n = 5) and (b) rehydrated (n = 6) state

Plasma copeptin concentrations remained elevated for a longer period of time post-biopsy during HYPO compared to RE, though there was also greater variance in individual responses (**Figure 3**). It took between 30 and 60 minutes for copeptin concentrations to return to baseline during HYPO, compared to < 30 minutes for all but two participants during RE (**Figure 3**). About half the participants had strong (more than doubling of basal levels) copeptin responses compared to the other half where less than doubling of copeptin concentrations typically occurred pre- to post-biopsy (with some participants barely responding to the biopsy during RE). The greater the initial copeptin biopsy response, the longer it took for copeptin to return to pre-biopsy levels (**Figure 3**).

**Figure 3.**
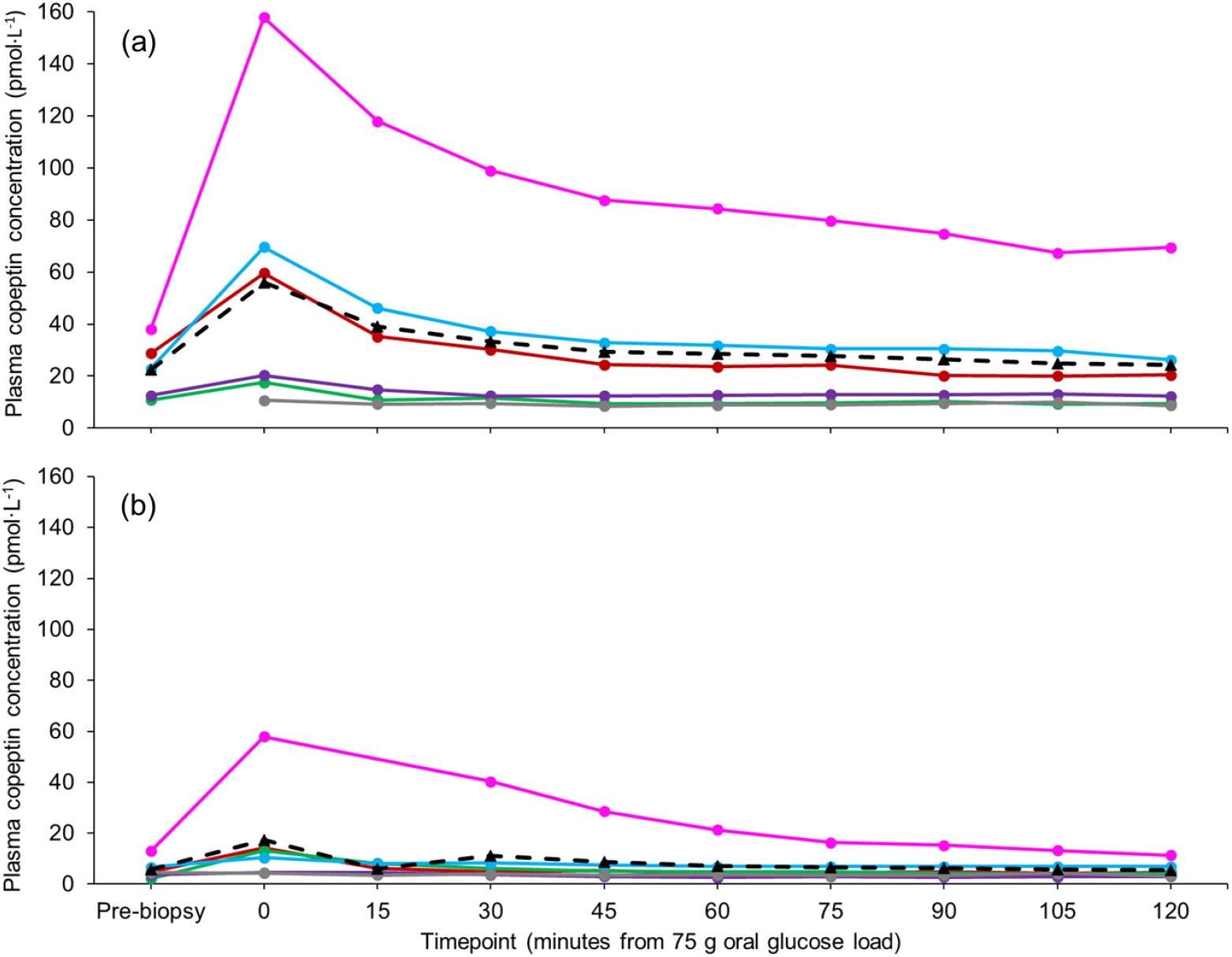
Individual (coloured lines) and average (black dashed lines) plasma copeptin responses pre- and post-biopsy and glucose ingestion the (a) hypohydrated and (b) rehydrated (n = 6) state

The level of total body water loss correlated strongly with the change in copeptin pre- to post-biopsy, explaining ∼57 % of variance; those at the highest level of hypohydration (assessed by percent body mass loss compared to baseline euhydration) tended to have a greater increase in copeptin after the biopsy (for each percent body mass retained, Δ copeptin was β −19.2, 95 % CI −42.7, 4.2 pmol·L^−1^ lower, *p* = 0.085), with no trend seen during RE (*p* = 0.974) (**Figure 4**).

**Figure 4.**
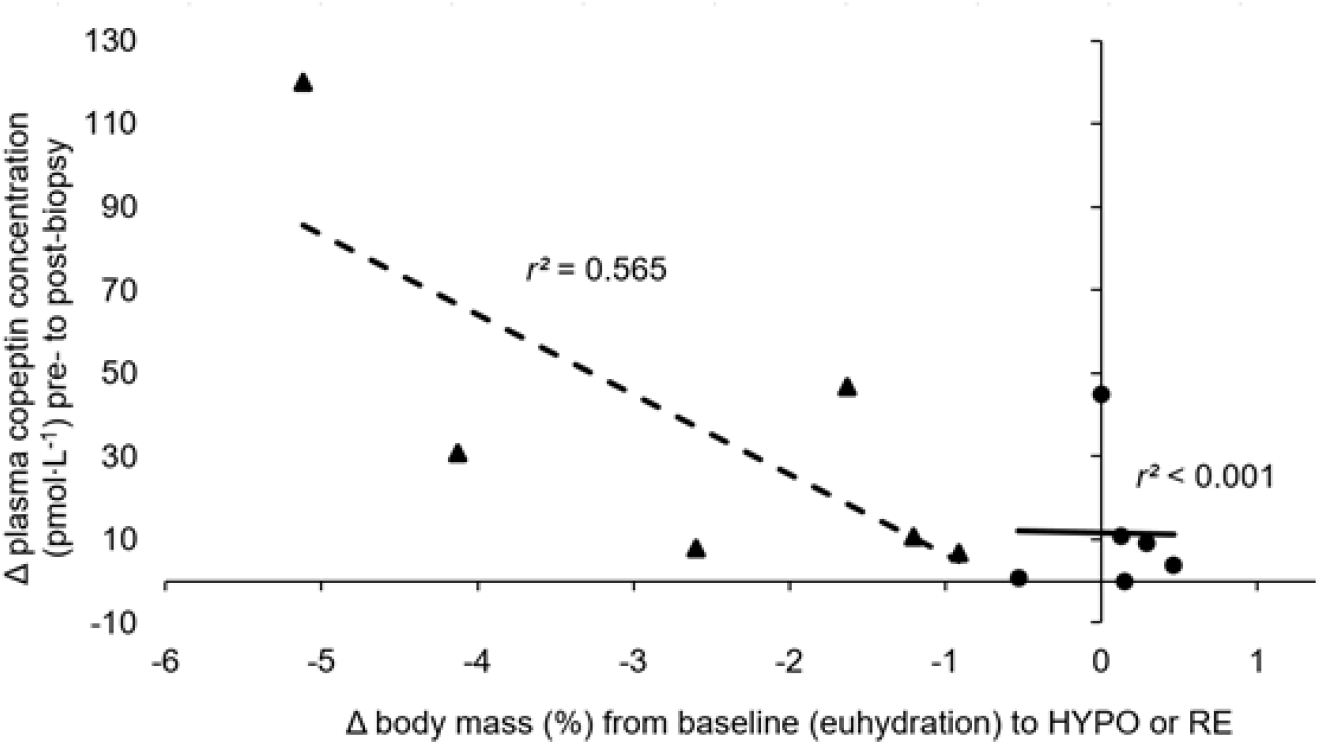
Percent body mass change from baseline euhydration to HYPO (triangles and dashed line) or RE (circles and filled line) and change in copeptin concentration pre-to post-biopsy (n = 6). Abbreviations: HYPO, hypohydration; RE, rehydration

As shown in **Figure 5**, copeptin, ACTH, and cortisol all responded to the muscle biopsy in both HYPO and RE. However, both ACTH and cortisol responded similarly regardless of hydration status, whereas copeptin had both a larger difference as well as a sharper peak during HYPO compared to RE. By ∼60 minutes post-OGTT, all three hormones were back to roughly baseline concentrations. The relationship between copeptin, ACTH, and cortisol pre- to post-biopsy is discussed further in the Supplementary Information.

**Figure 5.**
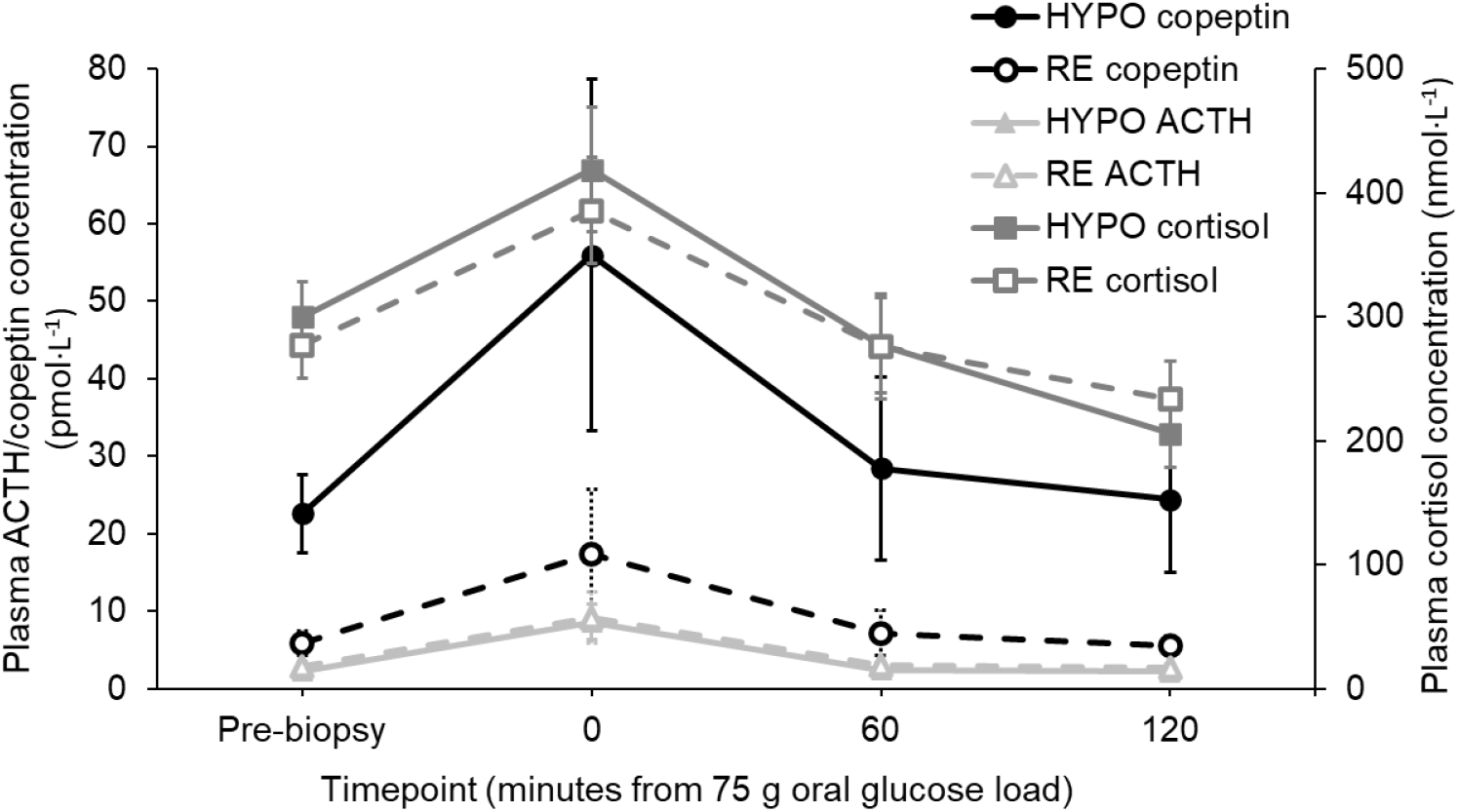
Time trend of copeptin, ACTH, and cortisol (n = 6). Error bars: standard error of the mean. Abbreviations: ACTH, adrenocorticotropic hormone; HYPO, hypohydration; RE, rehydration

## Discussion

In our original study, we included an opt-in muscle biopsy, thus unintentionally investigating the copeptin response to an acute physically stressful stimulus (Carroll *et al*., 2019). In this sub-analysis of the six participants who opted in for the muscle biopsies, after an acute bout of hypohydration at approximately 2.5 % average body mass loss, no notable changes were found in cardiometabolic metabolite or hormone concentrations. This is despite a meaningful increase in plasma copeptin and serum osmolality along with a ∼3.3 % reduction in cross-sectional muscle area, measured as a proxy for cell volume via pQCT. Overall, there was a consistent post-biopsy spike in copeptin which had returned to pre-biopsy levels within 60 minutes of instigating the OGTT (and in most participants, within 15-30 min). The post-biopsy spike in copeptin was higher and more prolonged during HYPO compared to RE; the magnitude of difference between HYPO and RE was only notably different immediately after the biopsy (timepoint 0 min), and 15 minutes post-OGTT in which there was a greater peak and slower return during HYPO.

The strong copeptin response suggests AVP, is a potent and acutely reactive stress hormone in humans which can be reflected in copeptin, independent of hydration status; an effect which to our knowledge has never been shown before. Further, hydration status can moderate this response, with hypohydration exacerbating and prolonging the copeptin response, as well as leading to higher absolute circulating copeptin concentrations. Previous research has shown an acute reduction in circulating plasma copeptin concentrations after a 1 L bolus of water (Enhörning *et al*., 2019). This reduction occurred within 30 minutes, somewhat corroborating our data in terms of time trends; due to the delay between finishing the biopsy and instigating the OGTT, our 15 minute time point is indicative of changes within ∼30 minutes of the biopsy ending.

Further, and notwithstanding the limitations of this secondary analysis including a small sample size, our data add confidence that copeptin can be used as a marker of acute stress, despite its longer half-life compared to AVP (Beglinger, Drew & Christ-Crain, 2017). Such information has applications in both research and clinical practice. Due to the instability of AVP, copeptin is increasingly being recognised as a valid surrogate marker which can aid research into acute stresses (such as hypohydration). Equally, copeptin is often measured clinically (such as being an early diagnostic tool for myocardial infarction; Keller *et al*., 2010). Our analyses offer assurances that high copeptin concentrations are unlikely to be delayed effects from acute interventions or stressful events and can be accurately used to assess current and acute stress. However, hydration status may be important to consider as even those with myocardial infarction exhibit lower copeptin concentrations than some of our participants who were hypohydrated (before the biopsy) (see also Carroll *et al*., 2019). Whilst the likelihood of this level of hypohydration is low in the general population, those presenting in hospital often have symptoms of hypohydration (e.g. Rowat, Graham & Dennis, 2012); thus hydration status should not be left unaccounted for when making clinical assessments based on copeptin.

Patients with acute myocardial infarction often have elevated copeptin concentrations and a cut-off of < 10 pmol·L^−1^ has been proposed to safely rule out myocardial infarction (Mockel, Searle, 2014). In the present analysis, median pre-biopsy copeptin concentrations were within this threshold when participants were well-hydrated (4.72 pmol·L^−1^), but far exceeded this cut-off when hypohydrated (22.83 pmol·L^−1^). Further, median post-biopsy (within ∼15 min) copeptin concentrations were above this cut-off when participants were both hydrated (13.17 pmol·L^−1^) and hypohydrated (59.65 pmol·L^−1^). Accordingly, when making clinical assessments based on copeptin values, hydration status needs to be taken account of, as do other forms of stress (such as deep wounds) which may exceed the 10 pmol·L^−1^ threshold without having clinical relevance. However, further research is needed to understand the interactions between hydration status and copeptin in a clinical setting considering our data represent healthy adults with no comorbidities or recent cardiac events.

Interestingly, despite a meaningful increase in plasma copeptin after HYPO, there were minimal differences between HYPO and RE for plasma ACTH and cortisol concentrations. This perhaps supports evidence that AVP alone is a weak secretagogue of ACTH; it is likely that the biopsy induced a CRH response which was the primary driver of the (small) ACTH response in this study. However, as AVP theoretically works synergistically with CRH on ACTH secretion (Gibbs, 1986, Miller, O’Callaghan, 2002, Goncharova, 2013), it is unexpected that there were no hydration status differences in ACTH or cortisol concentrations, considering the markedly and consistently higher plasma copeptin during HYPO. This suggests that our findings are the result of the two separate AVP neural pathways. Firstly, AVP was (basally) high during HYPO likely due to activation of hypothalamic magnocellular neurons in the paraventricular nucleus and supraoptic nucleus (Goncharova, 2013). Secondly, AVP was likely produced as part of the stress response in parvocellular neurons of the paraventricular nucleus, which produces CRH and regulate the HPA axis (Goncharova, 2013).

However, this explanation leaves some AVP (copeptin) unaccounted for in the data, since HYPO induced greater net change in copeptin post-biopsy relative to RE. It is unclear why magnocellular neurons would produce more AVP in response to stress, though this cannot be ruled out as a possibility. It seems more likely that parvocellular neurons produced the excess AVP during HYPO, though this fails to explain why there were no ACTH or cortisol differences between HYPO and RE. One preposition may be a counterregulatory reduction in CRH; whether and if this is an explanation should be further explored, including why this might occur with hypohydration. It is of course noted that we measured ACTH and cortisol hourly (compared to 15-minutely for copeptin), perhaps missing acute and rapid trends.

After the biopsy, plasma copeptin and serum glucose concentrations increased with HYPO (though the increase in glucose was not statistically different compared to the increase in RE). One hypothesised mechanism for AVP leading to higher glycaemia is via the HPA axis; thus, it is interesting that equivalent increases in ACTH and cortisol were found during both HYPO and RE conditions. If this AVP-HPA model of glycaemic dysregulation was correct, we would expect to see the increase in plasma glucose concentrations during HYPO correspond with the spike in cortisol. As such, considering cortisol was similar between trial arms, we would expect that glucose concentrations be similar between the two trials.

The lack of this response adds credence to recent hypotheses that this AVP-glucoregulatory model may not explain previous hydration and glycaemia observations (Carroll, James, 2019); rather supraphysiological concentrations may be needed to cause a small increase in glycaemia (which we may have seen post-biopsy with HYPO), likely mediated by V1 receptor-induced hepatic glycogenolysis (Koshimizu *et al*., 2012, Spruce *et al*., 1985). Indeed, previous research in those with acute myocardial infarction (without known diabetes) has indicated that copeptin reflects stress but is not implicated in the pathogenicity of glucose dysregulation (Smaradottir *et al*., 2017). In other words, those who are more susceptible to stress are at higher risk of glucose dysregulation, as well as having a stronger AVP response.

Those who lost the greatest amount of body mass during HYPO had the highest change in their pre- to post-biopsy copeptin concentration. No relationship was seen after RE, possibly due to there being a lower variance of body mass change, and greater weight stability from euhydration to RE; in other words, it is unknown whether there would be a lower copeptin response if weight was gained through hyperhydration. Speculatively, this may suggest that those more prone to rapid hypohydration are also more prone to greater AVP responses to acute physical stress, perhaps to maximise the bodies defence against hypohydration in fight-or-flight situations.

Notably, two participants with the strongest copeptin response to the biopsy were also highly physically active (these participants burnt approximately nearly double the energy of most other participants in physical activity based on their pre-trial physical activity data; data available: https://doi.org/10.15125/BATH-00547). Sweat responses are known to occur earlier in those with greater fitness (Ichinose-Kuwahara *et al*., 2010); as such perhaps an early and strong copeptin stress response is a sign of physical fitness, potentially offering insight into clinical interpretation of these values. Whilst no inferences can be made with the present data, this offers a hypothesis for further research to investigate the role of physical fitness in copeptin stress responses.

Additionally, those who had the strongest copeptin response to the biopsy, also had the strongest copeptin response to HYPO. Previous research has shown copeptin non-response to psychological stress in those with diabetes insipidus (Siegenthaler *et al*., 2014); thus taken together it may be that there are distinct copeptin stress strong- and weak-responders. Such a notion is supported by Enhörning *et al*. (2019) who found copeptin responders and non-responders to a week-long intervention increasing water intake, demonstrating an inverse responders/non-responders trend (i.e. increasing water, rather than HYPO In the present study). This secondary analysis, nor the primary study, were equipped to give reliable insights on this; as such this is something worthy of future research.

This paper adds to the limited current understanding of the human AVP response to acute stress under different hydration states, however many limitations are present within this sub-analysis, meaning any inferences made should be taken cautiously and exploratively. Firstly, this is an exploratory analysis of a randomised crossover trial investigating the effect of hydration status on glycaemia; thus inducing a stress response was an unintended consequence of the original study and not the primary objective. Secondly, our analysis included a minority subsample of mostly males from the original study, reducing the power to detect differences and reducing the generalisability to women who may exhibit differences in copeptin and/or the stress response (Herman *et al*., 2016). Thirdly, acute bouts of hypohydration may not reflect daily fluctuations in fluid balance which may be more reflective of clinical cases where copeptin is measured. Fourthly, our time-trend data come with the addition of feeding (75 g glucose); previous work has suggested a hydration state-feeding-cortisol interaction in those with type 2 diabetes withdrawn from medication (Johnson *et al*., 2017). Whilst there was no evidence of a similar interaction in our study, this cannot be ruled out without further investigation.

Additionally, we used Lidocaine as a local anaesthetic, which may have influenced the stress response in that physical pain was minimised; this may indicate that psychological stress impacted AVP more acutely. Since AVP is a regulator of psychological stress (e.g. depression; Landgrad, 2006), there may be a psychological stress-hydration status interaction for copeptin which does not translate to other stress hormones (in this case ACTH and cortisol, which both responded to stress but not differentially according to hydration status—perhaps the addition of physical pain would have differentiated these responses by hydration status). However, psychological stress only appears to induce a small (< 2 pmol·L^−1^) increase in copeptin concentrations (Siegenthaler *et al*., 2014), which is below what we found, even under rehydrated conditions (11.59 pmol·L^−1^ increase).

Nonetheless, the original study was, to our knowledge, the most tightly controlled hydration and health study in the literature, with both compliance to the protocol and hydration status confirmed by several measures. Further, there are several ethical issues with inducing stress, so these data have provided a unique opportunity to investigate the copeptin stress response in humans in different hydration states. Thus we hope this exploratory analysis and discussion will help generate new hypotheses.

Overall, these data highlight there is great individual variance regarding how copeptin responds to acute stress, with the general trend being that hypohydration exaggerates and prolongs the copeptin stress response. We have provided highly controlled evidence that copeptin responds dynamically and is therefore likely suitable to be used to understand acute stress. However, contrary to expectations, the corresponding ACTH and cortisol responses were not as predicted, bearing little relation to hydration-related copeptin responses, perhaps due to synthesis via different regulatory pathways in the brain. Researchers should consider their study design when utilising stressful techniques such as biopsies as some participants may have extended copeptin responses which may skew other measures.

## Declarations

No funding was received for conducting this study. HAC has received research funding from the Economic and Social Research Council, the European Hydration Institute, and the Esther Olssons stiftelse II & Anna Jönssons Minnesfond; has conducted research for Tate & Lyle; and has received speakers fees from Danone Nutricia Research. HAC and O.M. has received a research grant and consultancy fee from Danone Research.

## Availability of data and material

Data are available openly via doi: 10.15125/BATH-00547

## Authors’ contributions

Conceptualisation: Olle Melander; Methodology: Olle Melander and Harriet Carroll; Formal analysis: Harriet Carroll; Writing – original draft preparation: Harriet Carroll; Writing – review and editing: Olle Melander and Harriet Carroll

