## Supplementary Information for "Copeptin response to acute physical stress during hypohydration: An exploratory secondary analysis"

**Figure S1** demonstrates the difference in the copeptin response to the muscle biopsies according to HYPO and RE, highlighting HYPO exaggerated the copeptin response in all but 1 participant. There was however, large variation in participant copeptin responses, though the magnitude of the response in either HYPO or RE was strongly correlated ($r$ = 0.908). Thus, the copeptin stress response during RE explains approximately 82 % of the variation in the copeptin stress response during HYPO (β 2.33, 95 % CI 0.84, 3.82, $p$ = 0.012).

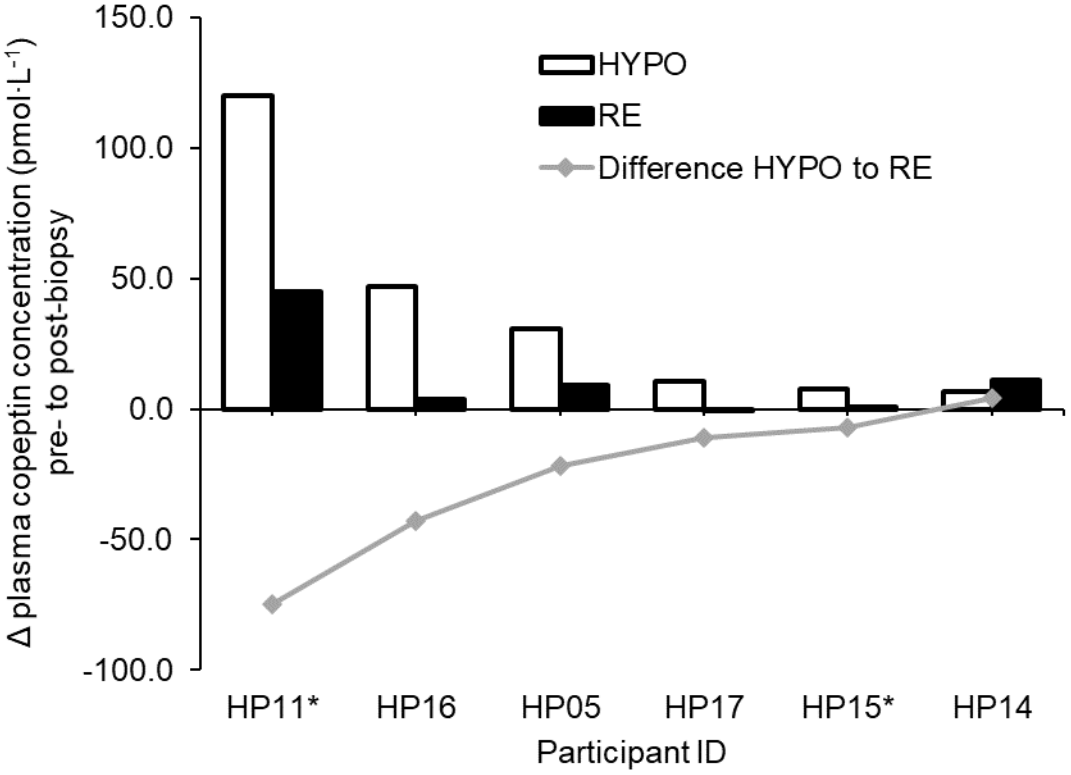


Figure S1. Change plasma copeptin concentration pre- to post-biopsy in HYPO and RE (n = 6). * Indicates participant was randomised to RE first. Abbreviations: HYPO, hypohydration; RE, rehydration

**Relationship between copeptin, ACTH, and cortisol pre- to post-biopsy**

The individual ACTH and cortisol responses to the biopsy are shown in Figures S2 and S3, respectively. Simple linear regressions were run to explore the relationship between pre- to post-biopsy change (Δ) copeptin and ACTH, and copeptin and cortisol. Due to non-linearity, the relationship between pre- to post-biopsy Δ in ACTH and cortisol was run with a quadratic term.

No significant relationship was found for pre- to post-biopsy Δ in copeptin *versus* ACTH after HYPO or RE, nor between Δ copeptin and cortisol during RE (Table S1, Figures S4a,b). After HYPO however, Δ in copeptin explained 67 % of variance in Δ cortisol, with each pmol·$L^{-1}$ increase in copeptin associated with a 0.10 nmol·$L^{-1}$ increase in cortisol (Table S1,

Figure S4b). Using a non-linear (quadratic) model, pre- to post-biopsy Δ ACTH explained 81 % of the variance in Δ cortisol during HYPO, and 96 % variance during RE (Table S1, Figure S4c).

Table S1. Relationship between pre- to post-biopsy Δ in copeptin, ACTH, and cortisol

| | $r^2$ | β | 95 % CI | *p*-value |
|---|---|---|---|---|
| | HYPO | | | |
| Δ copeptin vs Δ ACTH | 0.010 | 0.81 | -10.12, 11.74 | 0.847 |
| Δ copeptin vs Δ cortisol | 0.686 | 0.10 | 0.02, 0.59 | 0.042 |
| Δ ACTH vs Δ cortisol[a] | 0.811 | 92.34 | 10.47, 174.22 | 0.037 |
| | RE | | | |
| Δ copeptin vs Δ ACTH | 0.111 | 0.14 | -0.40, 0.67 | 0.518 |
| Δ copeptin vs Δ cortisol | 0.377 | 2.57 | -2.02, 7.16 | 0.195 |
| Δ ACTH vs Δ cortisol[a] | 0.961 | 33.35 | 20.00, 46.72 | 0.004 |

[a]Quadratic term used to account for non-linearity of data. Quadratic terms were significant for both models (HYPO $p = 0.040$; RE $p = 0.007$). Abbreviations: ACTH, adrenocorticotropin releasing hormone; CI, confidence interval; HYPO, hydration trial arm; RE, rehydration trial arm

These associations support that AVP (copeptin) alone is a weak secretagogue of ACTH, though the relationship appeared to be less weak during RE (i.e. when basal copeptin was lower). There was a stronger relationship between change in plasma copeptin concentration and change in cortisol, with this relationship being more pronounced during HYPO (i.e. when basal copeptin is higher). These results to our knowledge are unexplainable by current theory; this study was not equipped to give insight into such a finding. Thus, speculatively, they may suggest that AVP acts more directly on cortisol secretion via other mechanisms beyond ACTH secretion.

The non-linearity of the pre- to post-biopsy Δ in ACTH and cortisol concentrations was also unexpected, demonstrating a strong inverted-U shape curve. During HYPO, the response appears more leptokurtic (i.e. higher and earlier peak in the relationship), compared to the more platykurtic relationship during RE. This may suggest a more rapid secretion of cortisol in response to ACTH during HYPO, or could be highlighting a feedback loop preventing a prolonged stress response (i.e. the higher the ACTH change, the quicker the feedback leading to a reduced cortisol secretion during HYPO) (Goncharova, 2013[1]). The non-linearity of the curve may also indicate the slower acting nature of cortisol relative to ACTH. Of course, these ideas are speculative and should be further investigated, particularly due to the limited sample size of 6 participants, which is likely to give rise to spurious relationships.

[1] Goncharova, N.D. 2013, "Stress Responsiveness of the Hypothalamic–Pituitary–Adrenal Axis: Age-Related Features of the Vasopressinergic Regulation", *Frontiers in Endocrinology*, vol. 4, pp. 26.

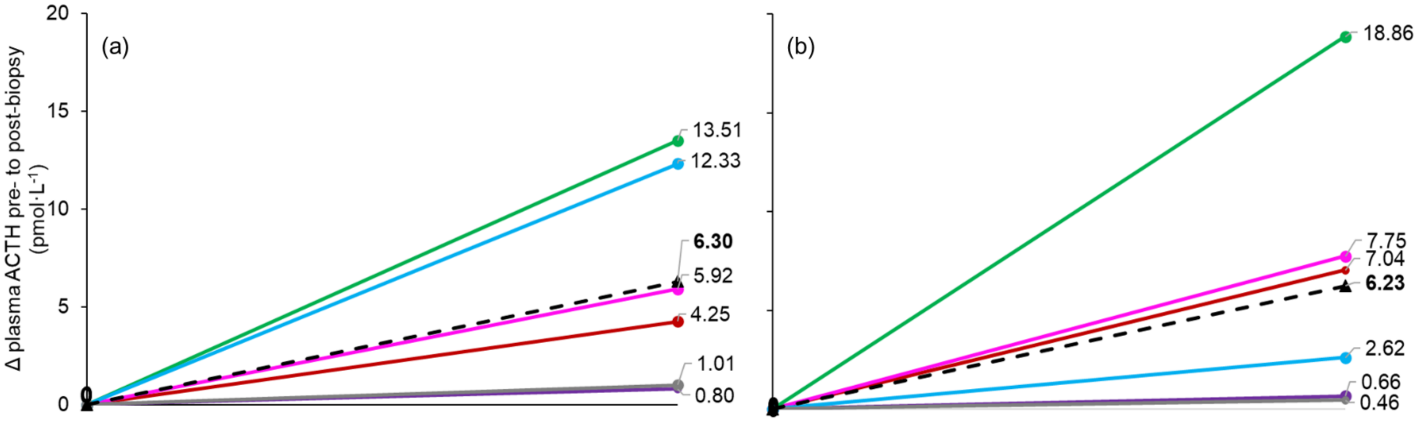


Figure S2. Individual (coloured lines) and average (black dashed lines) Δ plasma ACTH concentration pre- to post-biopsy in the (a) hypohydrated and (b) rehydrated state (n = 6). Abbreviation: ACTH, adrenocorticotrophic hormone

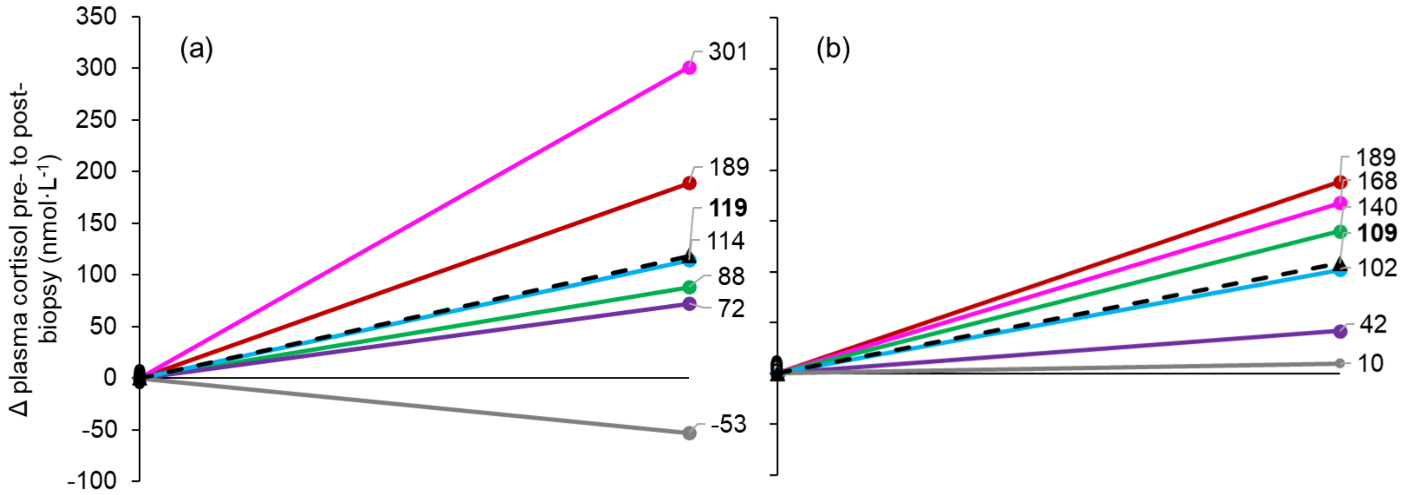


Figure S3. Individual (coloured lines) and average (black dashed lines) Δ plasma cortisol concentration pre- to post-biopsy in the (a) hypohydrated and (b) rehydrated state (n = 6)

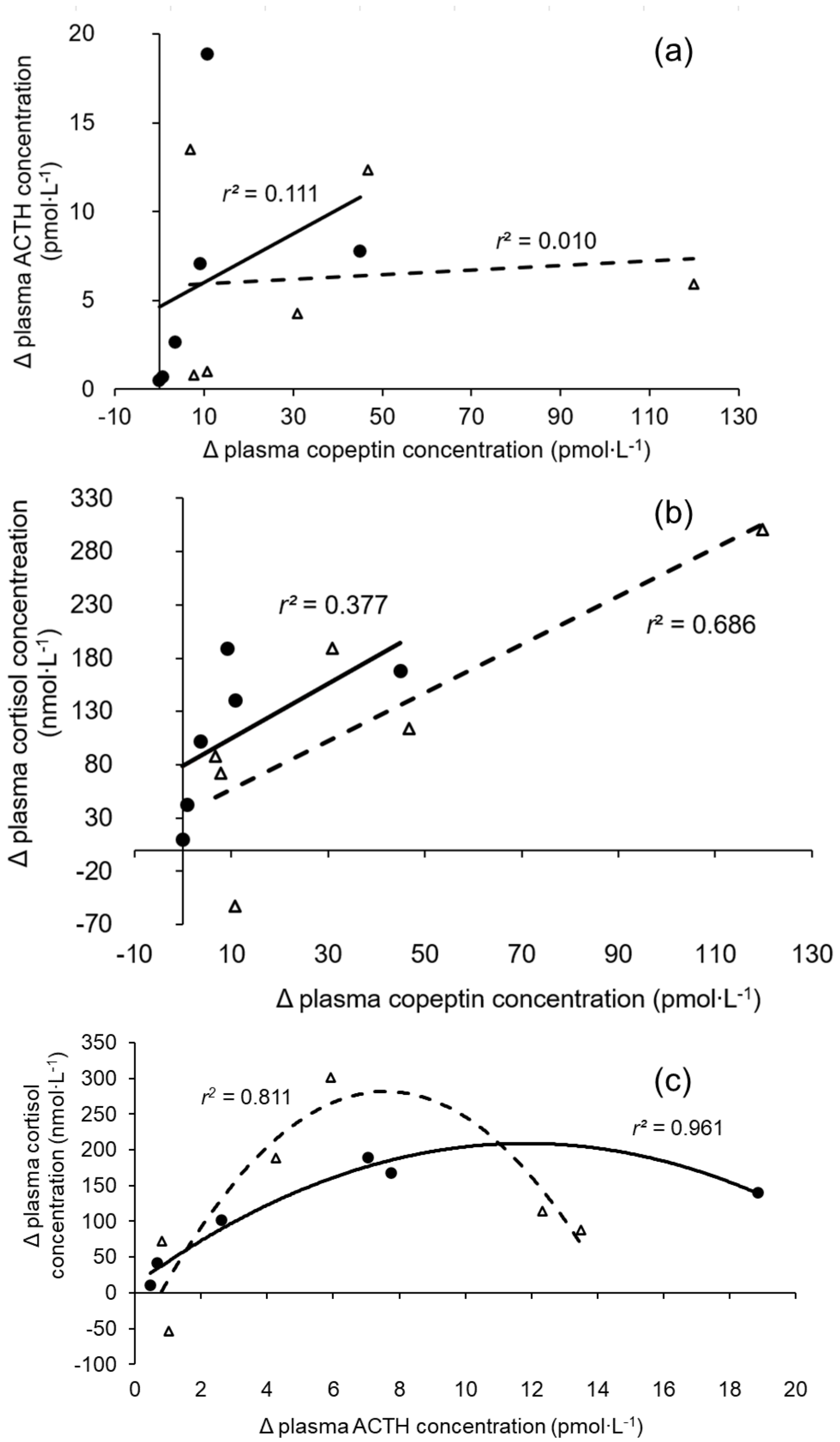


Figure S4. Relationship between the (a) change in plasma copeptin and ACTH, (b) change in plasma copeptin and cortisol concentrations, and (c) change in ACTH and cortisol pre- to post-biopsy during HYPO (triangles and dashed lines) and RE (circles and filled lines). Abbreviations: ACTH, adrenocorticotropin hormone; HYPO, hydration; RE, rehydration
